# Evolutionary Systems Biology Identifies Genetic Trade-offs In Rice Defense Against Above- and Belowground Attackers

**DOI:** 10.1101/2024.05.04.592539

**Authors:** Taryn S. Dunivant, Damaris Godinez-Vidal, Craig Perkins, Madelyn G. Lee, Matthew Ta, Simon C. Groen

## Abstract

Like other plants, wild and domesticated rice species (*Oryza nivara, O. rufipogon*, and *O. sativa*) evolve in environments with various biotic and abiotic stresses that fluctuate in intensity through space and time. Microbial pathogens and invertebrate herbivores such as plant-parasitic nematodes and caterpillars show geographical and temporal variation in activity patterns and may respond differently to certain plant defensive mechanisms. As such, plant interactions with multiple community members may result in conflicting selection pressures on genetic polymorphisms. Here, through assays with different above- and belowground herbivores, the fall armyworm (*Spodoptera frugiperda*) and the southern root-knot nematode (*Meloidogyne incognita*), respectively, and comparison with rice responses to microbial pathogens, we identify potential genetic trade-offs at the *KSL8* and *MG1* loci on chromosome 11. *KSL8* encodes the first committed step towards biosynthesis of either stemarane- or stemodane-type diterpenoids through the japonica (*KSL8-jap*) or indica (*KSL8-ind*) allele. Knocking out *KSL8-jap* and *CPS4*, encoding an enzyme that acts upstream in diterpenoid synthesis, in japonica rice cultivars increased resistance to *S. frugiperda* and decreased resistance to *M. incognita*. Furthermore, *MG1* resides in a haplotype that provided resistance to *M. incognita*, while alternative haplotypes are involved in mediating resistance to the rice blast fungus *Magnaporthe oryzae* and other pests and pathogens. Finally, *KSL8* and *MG1* alleles are located within trans-species haplotypes and may be evolving under long-term balancing selection. Our data are consistent with a hypothesis that polymorphisms at *KSL8* and *MG1* may be maintained through complex and diffuse community interactions.

## Introduction

Rice is consumed by approximately half the global population and is an important staple crop for securing the global food supply (Wing et al., 2018). Yet, elite rice varieties are profoundly vulnerable to both pathogens and invertebrate herbivore pests. As it stands, rice is more sensitive to such attackers than other staple crops (Savary et al., 2019). Moreover, these negative agronomic impacts of pests and pathogens are likely to deteriorate with the increased frequency of weather extremes and anticipated decreases in water availability associated with climate change (Dutta and Phani, 2023; Tonnang et al., 2022; Wang et al., 2022). This concern has become even more critical with the global push for direct-seeded rice cultivation. Direct-seeded rice cultivation reduces the amount of water, labor, and greenhouse gas emissions from current rice production methods. However, it will not be possible without resistance to pests and pathogens. Foundational research of molecular resistance mechanisms is essential for developing tools and novel approaches for pest and pathogen management in rice production.

Our approach is to harness the natural defense strategies that wild and domesticated rice evolved for crop improvement. We use the extensive genetic diversity of rice to discover resistance to biotic stressors, in particular herbivorous pests. Traditional rice cultivars have locally adapted to diverse agroecosystems (*e*.*g*. irrigated and rainfed lowland, deepwater, upland). Rainfed cultivars generally evolved more potent stress resistance mechanisms than irrigated cultivars as many stressors tend to be more prevalent agents of selection in rainfed agroecosystems. Furthermore, rice cultivars are classified by varietal groups grounded in the evolutionary history of domesticated rice and their wild relatives (temperate japonica, tropical japonica, *circum*-basmati, *circum*-aus, and indica). Temperate japonica is least similar to the other varietal groups in terms of cultivation environment. Temperate japonica is typically only grown in irrigated lowland systems at higher latitudes, whereas the other varietal groups are cultivated in a range of agroecosystems, often in (sub)tropical regions with more stable and intense herbivore activity. We can use these differences to compare the other varietal groups to temperate japonica. Other groups are more likely to have evolved potent anti-herbivore resistance mechanisms. This presents an opportunity to use a comparative framework for investigation. Differences in resistance between rice cultivars can be exploited to reveal the genetic basis of rice responses to herbivorous pests.

Revealing regulatory mechanisms of complex traits such as anti-herbivore resistance has the potential to provide insight into major biological properties of plants that promote translational research (Jung et al., 2008; Mochida and Shinozaki, 2011). For example, we can determine suites of genes that are functioning in tandem under specific conditions to achieve a precise phenotype. Furthermore, regulatory networks provide measurements of the plasticity of molecular mechanisms under fluctuating environmental stressors (Rivera et al., 2021). Thus, we are using evolutionary systems biology to identify molecular resistance mechanisms to herbivory in rice.

Plants are susceptible to above- and belowground herbivory on shoots and roots, respectively. This presents an opportunity for diverging selection pressures on rice defenses from one herbivore to another. Plant-parasitic root-knot nematodes (RKNs) of the genus *Meloidogyne* are important belowground herbivores in tropical rice agroecosystems (Dutta and Phani, 2023).

RKNs are obligatory sedentary endoparasites and cause characteristic root swellings, commonly called knots or galls, that serve as their feeding sites. In rice production, this can cause patchy fields and severe yield loss (Mantelin et al., 2017). Temperate japonica cultivars tend to be more susceptible to RKNs than cultivars from other varietal groups (Dimkpa et al., 2016; Zhan et al., 2018). Nematode resistance is lacking in many crops and advanced genetic tools could be a significant gain (Sikora et al., 2023).

An emerging aboveground herbivore found in rice agroecosystems is *Spodoptera frugiperda* or the fall armyworm (FAW). Its larvae are caterpillars that act as leaf-chewing herbivores and these have an affinity for grass crops. Levels of *S. frugiperda* infestation are linked linearly to increases in rice defoliation, reductions in plant and panicle densities, and reductions in rice yields (Pantoja et al., 1986). Temperate japonica cultivars also tend to be more susceptible to FAW and other chewing insect herbivores than cultivars from other rice varietal groups (Heinrichs, 1986; Wang et al., 2022). FAW is present year-round in tropical regions, but its presence elsewhere is restricted by coldest annual temperature (Early et al., 2018).

In tropical rainfed lowland rice environments RKN and FAW infestations often occur after standing water has evaporated from paddies during periods of drought, whereas rainfed upland environments are natural habitats for RKNs and further experience increased outbreaks of FAW when periods of drought are followed by rains (Bridge et al., 2005; Rose et al., 2000).

Rice is equipped with molecular defense mechanisms against herbivory. The ability of plants to launch a defense response often relies on recognition of an attacker. An important recognition mechanism is through intracellular nucleotide-binding leucine rich-repeat (NLR) immune receptors that evolve under species-specific selection. Following recognition of an attacker signaling cascades are induced that regulate defense responses such as phytoalexin accumulation, cell wall restructuring, deposition of waxes and suberin, production of reactive oxygen species, and protein reinforcements (Bernaola et al., 2021; Groen et al., 2016; as reviewed in Qi et al., 2018). A major line of defense in both roots and shoots is phytoalexin accumulation. Rice produces two major types of phytoalexins, phenolics (e.g., flavonoids, phenylamides) and diterpenoids (e.g., momilactones, oryzalexins, phytocassanes). Interestingly, diterpenoid synthase genes are known to be upregulated in response to various stressors, including herbivory (Schmelz et al., 2014), and rice displays inter-cultivar variation in the amount and diversity of diterpenoid molecules produced (Gu et al., 2019; Kariya et al., 2024, 2023, 2020, 2019).

Here, we leverage leaf and root gene co-expression networks constructed from a rice diversity panel that consisted of aus, indica, and japonica cultivars from irrigated and rainfed agro-ecosystems grown in wet and dry field conditions where plants were naturally subjected to biotic factors that included leaf-chewing herbivores and RKNs (Groen et al., 2022; Sandhu et al., 2015; Xu et al., 2021). Combining these networks with existing genome-wide gene expression datasets from plants that were experimentally subjected to individual attacks by FAWs and RKNs we identify modules of co-expressed genes that are enriched for herbivore-responsive genes and functionally test candidate genes for their role in defense against above- and belowground herbivores.

## Results

### Evolutionary systems biology identifies diterpenoids as anti-herbivore defenses in rice

We identified herbivore-responsive genes (Zhou et al. [2020], *M. incognita* [Fig.1; Supplementary Table S1,S2]; Kyndt et al. [2012], *M. graminicola* [Fig. S1; Supplementary Table S2,S3]; Leclerc et al. [2024], *S. frugiperda* [Fig. 2; Supplementary Table S4,S5]; Venu et al. [2010], *S. exigua* [Fig. S2; Supplementary Table S5,S6]) that significantly overlapped with fitness-linked modules of a gene co-expression network constructed from a rice diversity panel grown in wet and dry environments (Groen et al., 2022).

**Figure 1.**
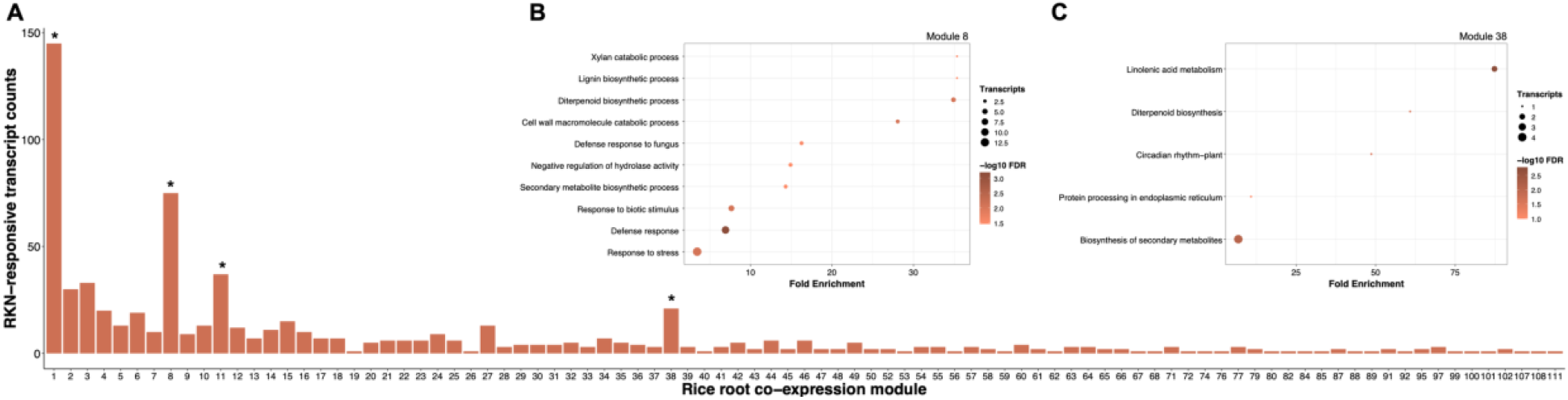
Distribution and gene set enrichment of root-knot nematode-responsive transcripts across the rice root co-expression network. A) Overlay of *Meloidogyne incognita*-induced transcripts at 6 days after infection on the rice root co-expression network. *, false discovery rate-adjusted p-value < 0.001 for a Fisher’s exact test on count data. B,C) Enrichment of defense-related processes as shown by Gene Ontology biological process enrichment analysis on root modules 8 and 38.

**Figure 2.**
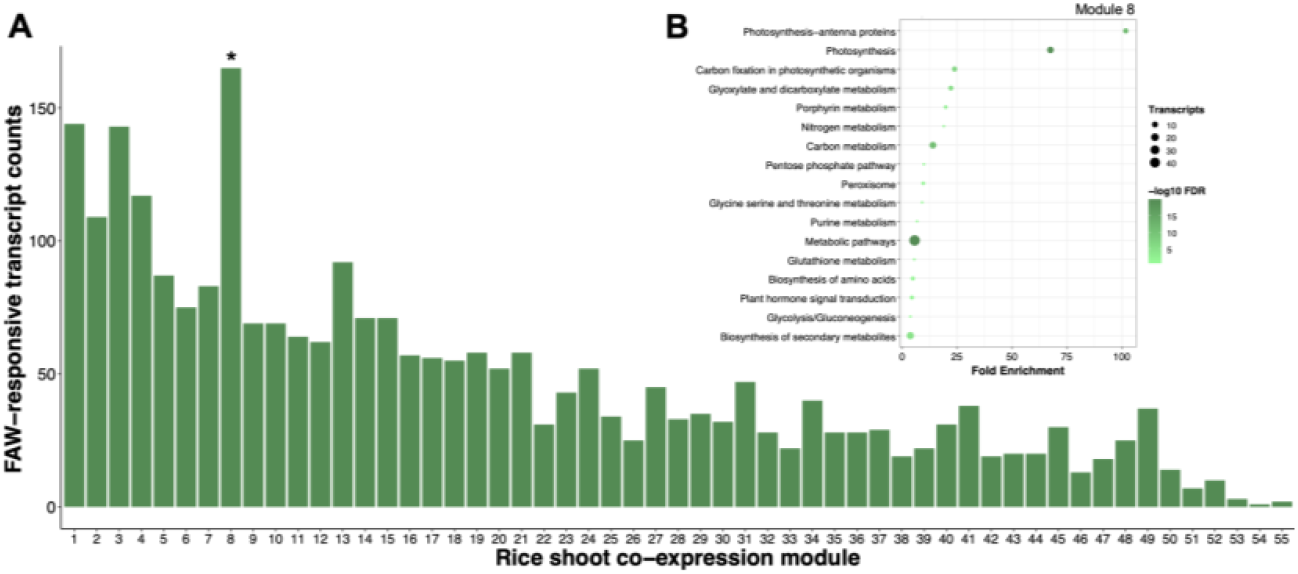
Distribution and gene set enrichment of fall armyworm-responsive transcripts across the rice shoot co-expression network. A) Overlay of *Spodoptera frugiperda*-induced transcripts at 2 hours after infestation on the rice shoot co-expression network. *, false discovery rate-adjusted p-value < 0.001 for a Fisher’s exact test on count data. B) Enrichment of defense-related processes as shown by the Gene Ontology biological process enrichment analysis of shoot module 8.

From our RKN analysis, we identified four modules from the root network that are enriched with RKN-responsive transcripts, specifically *M. incognita* transcripts (Fig. 1A). Of the total 971 differentially expressed genes reported, we find 145 in module 1 (p = 1.97x10^-4^), 75 in module 8 (p = 2.46x10^-14^), 37 in module 11 (p = 5.86x10^-5^), and 21 in module 38 (p = 3.39x10^-5^). For each of these gene sets we performed gene set enrichment analysis (GSEA) to categorize any gene ontology (GO) biological processes or Kyoto Encyclopedia of Genes and Genomes (KEGG) pathways of significance. Enrichment of modules 8 and 38 included defense-related processes and pathways, including ones related to cell wall restructuring, biosynthesis of pathogenesis-related proteins, stress signaling, and secondary metabolite synthesis (Fig. 1B,C; Supplementary Table S7,S8). Strongly represented among the genes related to secondary metabolites are those involved in diterpenoid metabolism (Os02g0569000, Os04g0179700, Os07g0190000, Os12g0491800, and Os06g0568600). The gene sets of modules 1 and 11 were not significant for any enrichment of processes or pathways.

From our FAW analysis, we identified one module from the shoot network that is enriched with armyworm-responsive transcripts (Fig. 2A). Of the 2,638 transcripts that were assigned to a module in the shoot network, only module 8 was enriched for armyworm-responsive transcripts (n = 165, p = 4.88x10^-12^). GSEA for this cluster showed enrichment for processes and pathways related to photosynthesis, generation of reactive oxygen species, hormone signal transduction, and secondary metabolite synthesis. Interestingly, shoot module 8 is enriched for genes involved in isopentenyl diphosphate (IPP) biosynthesis (Fig. 2B; Supplementary Table S9,S10; Groen et al., 2022). IPP and its isomer DMAPP are isoprenoid precursors, which in turn can form building blocks for diterpenoid biosynthesis (Murphy and Zerbe, 2020).

These findings suggest a role for diterpenoid metabolism as having potential roles in anti-herbivore defenses in both below- and aboveground tissues. We therefore further investigated the expression patterns of genes known to be involved in the biosynthesis of diterpenoid phytoalexins.

### Expression of diterpenoid metabolism genes responds to herbivory

To further investigate expression of genes underlying diterpenoid metabolism we explored the complete gene sets of root co-expression modules 8 and 38 (Groen et al., 2022). We discovered that a minimum of 15 diterpene synthase genes are within these two root modules and include ones encoding enzymes such as CPSs, KSLs, and CYP450s as well as all major branches that lead to the four classes of rice diterpenoids: oryzalexins (including oryzalexin S), phytocassanes, oryzalides, and momilactones (Fig. 3). This evidence suggests diterpenoid metabolism genes as being co-expressed in rice roots.

**Figure 3.**
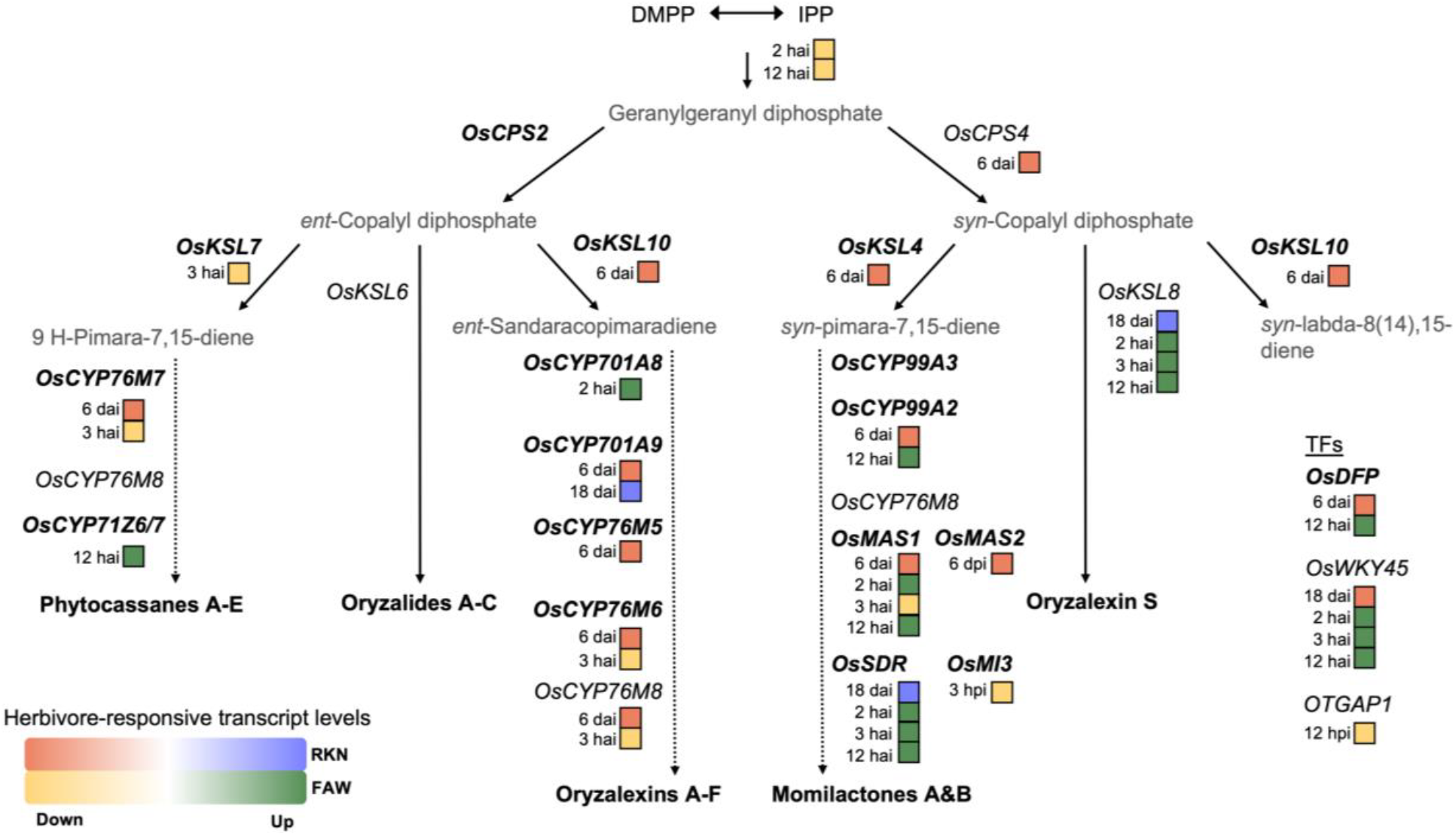
Expression analysis of genes underlying the diterpenoid biosynthetic pathway highlights co-expressed transcripts. Genes in bold belong to the same rice root co-expression module (module 8; Groen et al., 2022). Integrated with the diterpenoid pathway are the relative expression levels of diterpene synthase genes in response to RKNs (Zhou et al., 2020) and FAWs (Leclerc et al., 2024; Xue et al., 2024). RKN-responsive transcript levels: blue is upregulated and orange is downregulated; dpi is days post-infection. FAW-responsive transcript levels, green is upregulated and yellow is downregulated; hpi is hours post-infestation.

The majority of genes responsible for the production of phytocassanes A-E, oryzalides A-C, and oryzalexins A-F were downregulated upon RKN or FAW herbivory (Fig. 3; Supplementary Table S11). For RKNs, the same was true for the genes involved in the production of momilactones, although expression of these genes was generally upregulated in response to FAW feeding. Interestingly, however, among all the KSLs, which represent different branching points in the diterpenoid production pathway, expression of *OsKSL8* was unique in being upregulated in response to both RKN and FAW herbivory (Fig. 3).

*KSL8* is located on chromosome 11 and two alleles are known to segregate at this locus, *KSL8-jap* and *KSL8-ind*, with the former being prevalent in temperate japonica and the wild relative and ancestor of all japonica rice cultivars, *O. rufipogon*, and the latter being prevalent in tropical japonica (through introgression) and cultivars belonging to the indica and aus sub-populations as well as their respective wild relative and ancestor, *O. nivara* (Kariya et al., 2024; Zhao et al., 2023). KSL8-jap and -ind both act downstream of CPS4 with the *KSL8-jap* allele encoding the first committed step in the synthesis of stemarane-type diterpenoids, including oryzalexin S, and the *KSL8-ind* allele encoding a stemod-13-ene synthase that does not engender oryzalexin S production and rather represents the committed step in the production of stemodane-type diterpenoids (Kariya et al., 2024; Zhao et al., 2023). We hypothesized that *KSL8* could partially underlie previously observed differences in the general level of anti-herbivore resistance of temperate and tropical rice cultivars (Dimkpa et al., 2016; Heinrichs, 1986; Wang et al., 2022; Zhan et al., 2018).

### KSL8-jap positively mediates defense against belowground herbivores

Based on these analyses, we tested if KSL8-jap could play a role in defending rice plants against herbivory by the RKN *M. incognita*. At three days after infestation (3DAI), *ksl8-jap* knockout plants had attracted significantly more second-stage juvenile nematodes (J2s) of *M. incognita* than wild-type and *cps4* knockout mutant plants of cv. Kitaake (χ^2^ tests, p ≤ 0.00640 and p = 0.218, respectively; Fig. 4A). These patterns persisted at 10DAI and once J2s had infected plants they progressed significantly faster through development in *ksl8-jap* knockout plants than in wild-type and *cps4* knockout plants of cv. Kitaake (χ^2^ tests, p ≤ 2.00x10^-5^ and p = 0.275, respectively; Fig. 4B).

**Figure 4.**
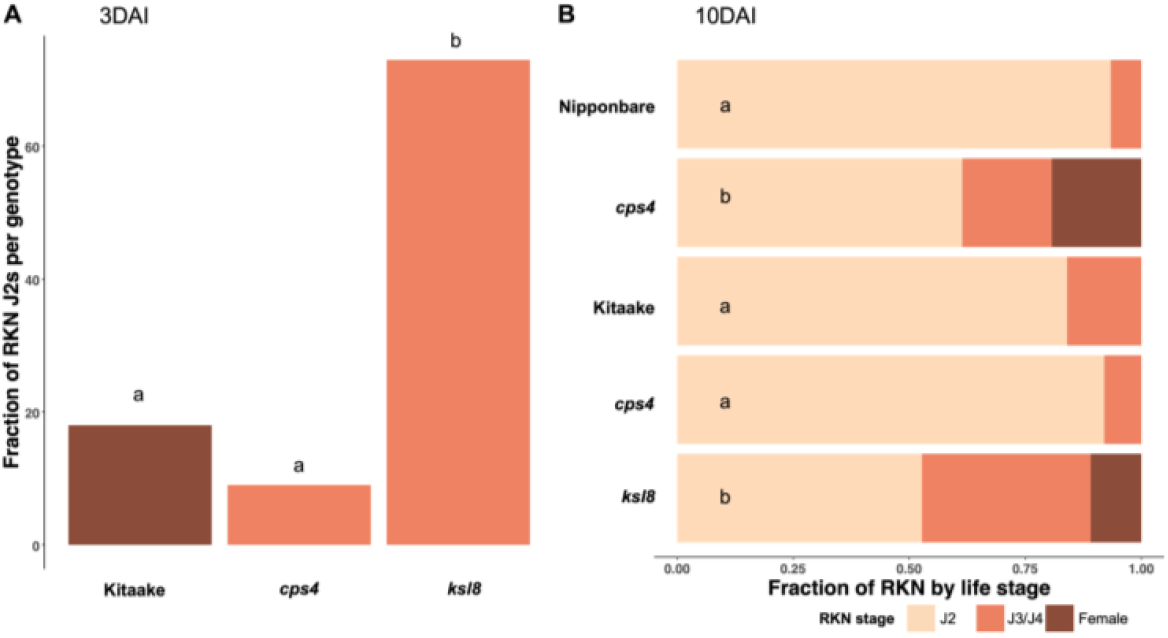
Changes in susceptibility of RKNs on *cps4* and *ksl8* knockout lines. A) Percent of total J2s able to infect root systems present in *cps4* and *ksl8* knockouts compared to their wild-type counterparts at 3 days after infection (3DAI). B) Fraction of RKNs by life stage in the root systems of *cps4* and *ksl8* knockouts compared to their wild-type counterparts at 10DAI.

Since previous work has shown enhanced susceptibility to RKNs of *cps4* knockout plants of cv. Kitaake at 14DAI (Desmedt et al., 2022), and since we did not observe significant differences at the earlier time-points of 3DAI and 10DAI, we decided to test a *cps4* knockout of cv. Nipponbare, which is known to accumulate higher diterpenoid levels than cv. Kitaake and contains the same allele at *KSL8, KSL8-jap* (Zhang et al., 2021). Indeed, at 10DAI *cps4* knockout plants of cv. Nipponbare showed significantly enhanced susceptibility to RKNs compared to wild-type plants (χ^2^ test, p = 1.13x10^-10^; Fig. 4B).

These results are in contrast with the hypothesis that allelic variation at *KSL8* may underpin observed general differences between temperate and tropical rice cultivars in their resistance to RKNs, since the temperate japonica allele *KSL8-jap* contributes to rice immunity. However, the effects of KSL8 can be overridden in the presence of the immune receptor MG1, which is segregating among tropical japonica, indica, and aus cultivars, but not temperate japonica cultivars (Dimkpa et al., 2016; Wang et al., 2023). This intracellular NLR immune receptor was previously shown to confer strong resistance to the RKNs *M. graminicola* and *M. javanica* upon recognition of nematode attack through a hypersensitive response (Lahari et al., 2020; Wang et al., 2023). We found that rice cultivars carrying *MG1* are strongly resistant to *M. incognita* as well, with nearly zero galling in cultivars LD24 and Khao Pahk Maw (two-tailed t-test relative to cv. Kitaake, p = 1.7 x10^-4^ and p = 4.23x10^-4^, respectively; Fig. 5A) along with the inability of juveniles to reach reproductive stages (in both comparisons: χ^2^test, p = 9.46x10^-48^; Fig. 5B). Perhaps coincidentally, at least one cultivar carrying *MG1*, Zhonghua 11, also accumulates much higher levels of diterpenoids than temperate japonica cultivars (Zhang et al., 2021).

**Figure 5.**
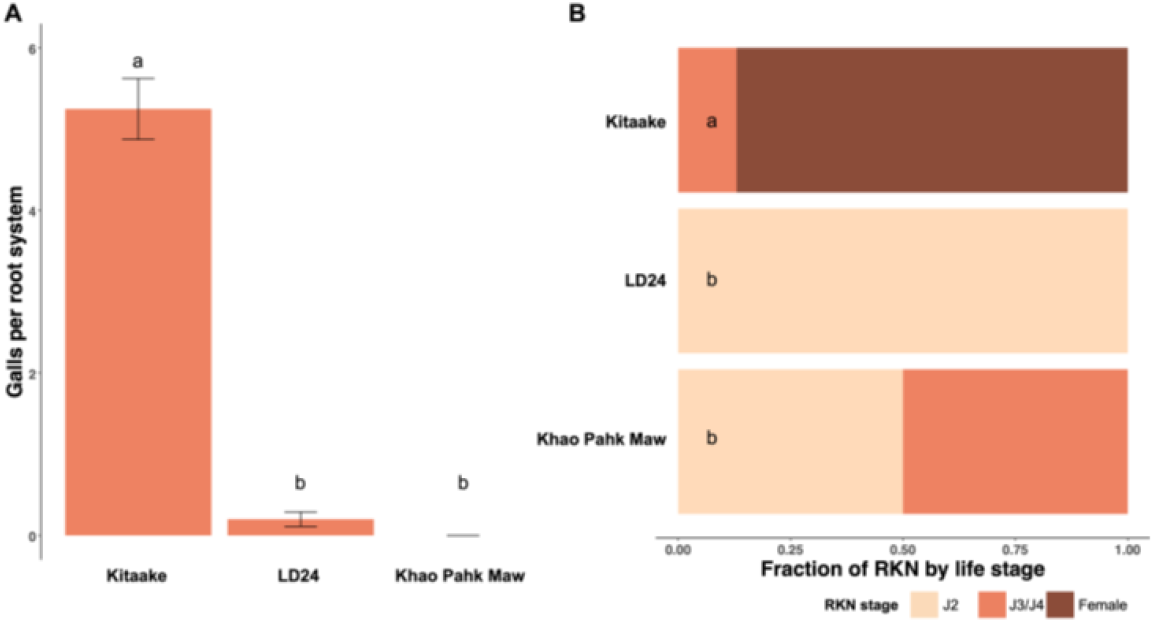
Effects of MG1 on infection rates of *M. incognita*. A) Average number of galls per root system for cv. Kitaake, LD24, and Khao Pahk Maw. B) Fraction of RKNs by life stage in the root systems of cv. Kitaake, LD24, and Khao Pahk Maw.

### KSL8-jap negatively mediates defense against aboveground herbivores

Next, we tested rice resistance to leaf feeding by FAW caterpillars. At 2DAI, compared to wild-type plants, *cps4* knockout plants of cv. Nipponbare showed enhanced resistance to feeding by second-instar FAW caterpillars with caterpillars gaining significantly less weight on leaves of the mutant (t-test, p = 0.0252; Fig. 6A). As for RKN herbivory, also for FAW herbivory the difference between *cps4* knockout plants and wild-type plants was less obvious for cv. Kitaake than for cv. Nipponbare, but the data did confirm a pattern of increased resistance to caterpillar feeding in the knockout line (one-tailed t-test, p = 0.0329; Fig. 6B).

**Figure 6.**
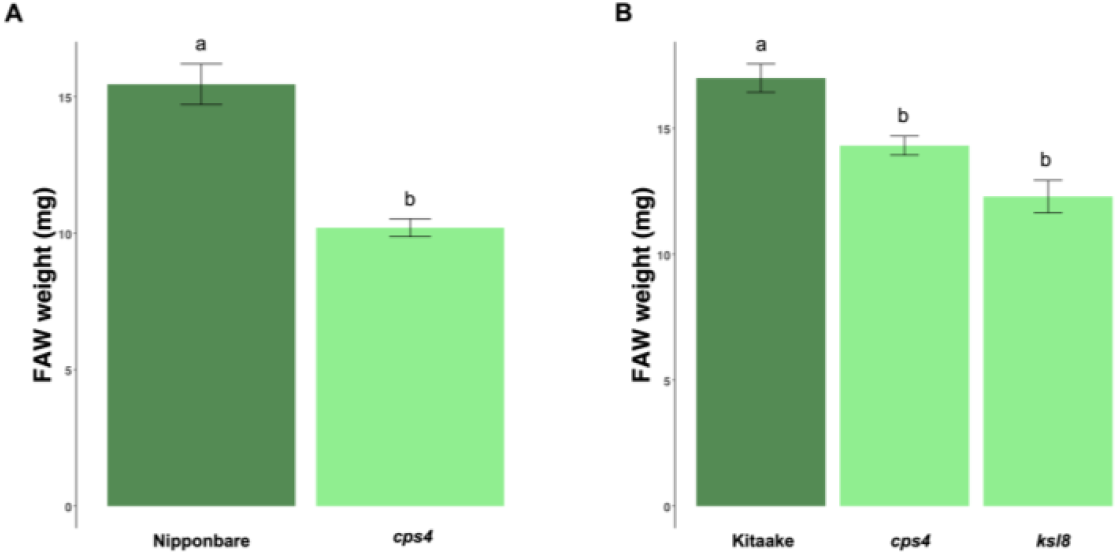
Changes in FAW weight from feeding on *cps4* and *ksl8* knockout lines. A) Weight of FAWs after feeding for two continuous days on *cps4* and wild-type cv. Nipponbare leaves. B) Weight of FAWS after feeding for two continuous days on *cps4, ksl8*, and wild-type cv. Kitaake leaves.

Like the *cps4* mutant, also the *ksl8-jap* mutant in cv. Kitaake showed an opposite pattern for resistance against FAWs compared to resistance against RKNs: caterpillars gained significantly less weight on leaves of *ksl8-jap* mutant plants than on those from wild-type plants (t-test, p = 0.0367; Fig. 6B). The results for the cv. Kitaake knockout mutants *cps4* and *ksl8-jap* were confirmed in a second experiment with FAW caterpillars gaining significantly less weight on leaves of either mutant compared to wild-type leaves (t-tests, p = 0.00401 for *cps4* and p = 0.0302 for *ksl8-jap*, respectively; Supplementary Fig. S3).

Since tropical rice cultivars from the tropical japonica, indica, and aus varietal groups are known to carry an alternative allele (*KSL8-ind*) at the *KSL8* locus that, similar to knocking out *KSL8-jap*, prevents production of stemarane-type diterpenoids such as oryzalexin S, we decided to test if tropical rice cultivars would be more resistant to FAW caterpillars than temperate cultivars, which was indeed the case (t-tests, p ≤ 0.0131 relative to cv. Kitaake and p ≤ 0.0887 relative to cv. Nipponbare; Fig. 7).

**Figure 7.**
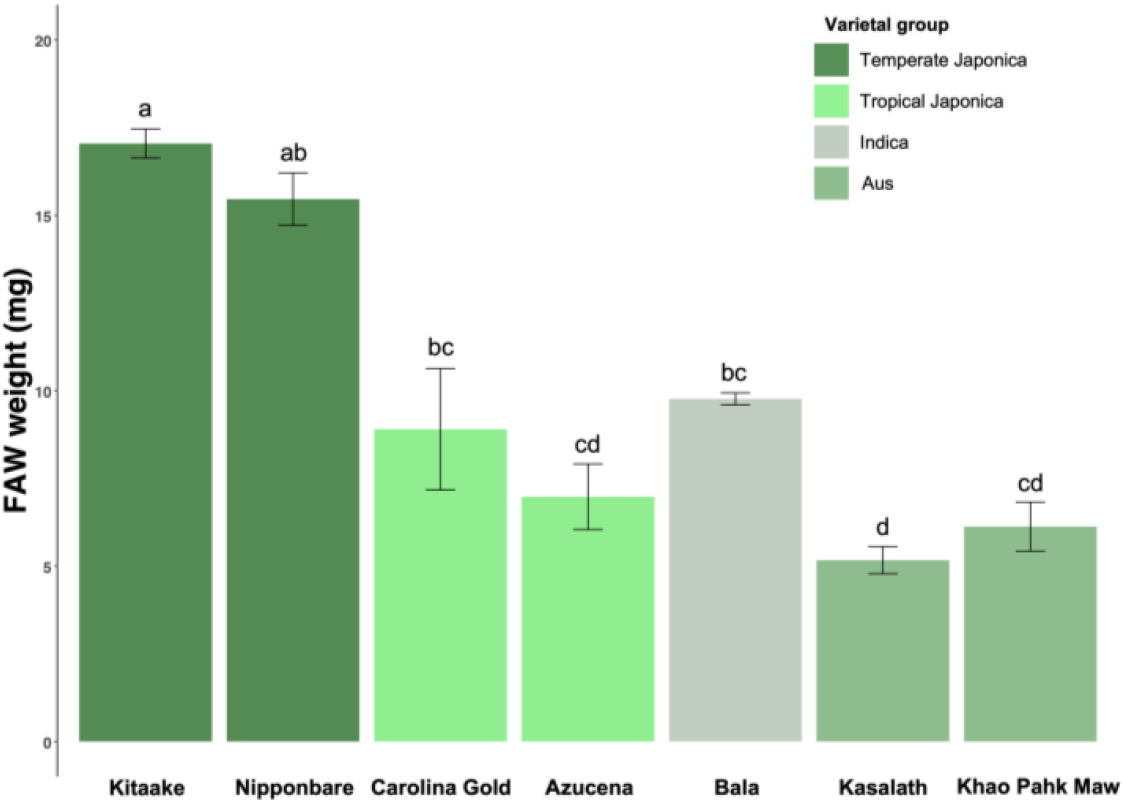
Effects of KSL8-ind and KSL8-jap on feeding rates of FAW. Weight of FAWs after feeding for two continuous days on cv. Kitaake, Nipponbare, Carolina Gold, Azucena, Bala, Kasalath, Khao Pahk Maw.

## Discussion

We identified potential anti-herbivore defense mechanisms in both above- and belowground rice tissues. Our GSEA for herbivore-responsive transcripts that are significantly enriched in gene co-expression modules linked with rice fitness in wet and dry environments revealed processes related to phytoalexin biosynthesis, cell wall restructuring, and immune responses. In this study, we investigated genes underlying production of phytoalexins, specifically diterpenoids, for their role in anti-herbivore defenses in rice.

It was evident from our analyses that genes underlying diterpenoid production are co-regulated. Of the estimated 23 genes involved in diterpenoid biosynthesis, 14 belong to a single root gene co-expression module (module 8), including the major regulator transcription factor *DITERPENOID PHYTOALEXIN FACTOR* (*DPF*). Furthermore, we know that two major gene clusters of diterpene synthases are located on chromosomes 2 and 4 (Miyamoto et al., 2016). Although expression of most of these genes is downregulated in response to RKNs and FAWs, expression of *KSL8* forms an exception, and diterpenoid biosynthesis is upregulated in response to several stressors and elicitors (e.g., Desmedt et al., 2022; Kato-Noguchi, 2009; Liu et al., 2024; Schmelz et al., 2014). For example, rice plants treated with exogenous phenylalanine ammonia-lyase inhibitor (AIP) display induced transcription of diterpene synthase genes and a high accumulation of phytocassanes, momilactones, and oryzalexins 3 days post-treatment. Interestingly, rice plants treated with AIP also show increased resistance to *M. graminicola* (Liu et al., 2024). Furthermore, a RKN-resistant mutant indica line showed upregulation of the diterpenoid synthesis pathway upon early recognition of *M. graminicola* (Dash et al., 2021). Previous studies have demonstrated diterpenoid production to be inducible by jasmonic acid (JA), a hormone that regulates anti-herbivore and anti-fungal defenses (Shimizu et al., 2013). RKNs and FAWs appear to avoid or prevent induction of JA-regulated immune responses because, although exogenous application of MeJA increases diterpenoid production and resistance to *M. graminicola* and *S. frugiperda* in rice (Nahar et al., 2011; Stout et al., 2009), expression of many genes underlying diterpenoid synthesis is downregulated upon herbivory. RKNs and potentially FAWs may avoid inducing most branches of diterpenoid metabolism.

Rice diterpenoids are diverse and have distinct functions (Li et al., 2021; Rayee et al., 2024). In our study, we found distinct and conflicting roles for the diterpenoid biosynthetic enzymes CPS4 and KSL8 in mediating defenses against RKN and FAW herbivory. A similar pattern has been observed for above- and belowground microbial pathogens. CPS4 has opposite effects on rice resistance to bacterial and fungal pathogens, functioning negatively in host resistance to the leaf bacterial pathogen *Xanthomonas oryzae* pv. *oryzae* and positively in non-host resistance to the root fungal pathogen *Magnaporthe poae* but not in host resistance to the related fungal pathogen *M. oryzae* (Lu et al., 2018). It was found previously that RKN herbivory increases for *M. graminicola* when *CPS4* is knocked out in cv. Kitaake (Desmedt et al., 2022). Our findings for cv. Kitaake infected by *M. incognita* showed a similar trend, but this was not significant. This may have been due to the earlier time points we chose to assess nematode numbers because we did identify a significantly enhanced susceptibility of *cps4* knockout plants of cv. Nipponbare, which is known to accumulate higher levels of diterpenoids than cv. Kitaake (Zhang et al., 2021). Furthermore, as was observed previously for *X. oryzae* pv. *oryzae*, we identified increased resistance to FAW in *cps4* knockouts. This opposing pattern of herbivore performance for RKNs and FAWs in *cps4* knockout lines was also visible for our *ksl8-jap* knockout line.

Temperate and tropical rice cultivars are known to segregate for a polymorphism in *OsKSL8* with temperate japonica cultivars possessing the *KSL8-jap* allele that encodes a stemar-13-ene synthase, which engenders production of oryzalexin S, and tropical japonica, indica, and aus cultivars carrying the *KSL8-ind* allele that encodes a stemod-13-ene synthase, which does not engender oryzalexin S production but rather represents the committed step in the production of diterpenoids of the stemodane family (Kariya et al., 2024; Zhao et al., 2023). Although some of the stemodane family diterpenoids possess antiviral activity (Morrone et al., 2006), it is unclear if they could affect invertebrate herbivores. While our results suggest they might, more experimental work will be needed to verify this. Our results further point to a potential role for oryzalexin S in rice defense against RKNs, but also this requires further experimental corroboration.

*Oryza sativa* ssp. japonica was domesticated from the (sub)tropical perennial *O. rufipogon* (Choi et al., 2017), which contains the *KSL8-jap* allele that contributes to anti-RKN defenses. It may be especially important for perennial herbaceous plants to protect their root systems from nematode herbivory since the roots persist in the soil for years. However, as japonica rice was domesticated and became an annual crop and as cultivation practices developed that avoid or resist prolonged exposure to active RKNs—for example through slash-and-burn agriculture, crop rotation, and flood irrigation—the increased RKN resistance that the *KSL8-jap* allele confers may have become less important. This could have facilitated the current situation in which the *KSL8-ind* allele has become the most prevalent allele in tropical japonica, indica, and aus cultivars.

Another factor may be that KSL8-mediated resistance to RKNs can be overridden in tropical rice cultivars by hypersensitive resistance through the immune receptor MG1. The *MG1* gene resides in an NLR gene-rich genome region on chromosome 11 that contains trans-species haplotypes (Zhou et al., 2021). Certain NLR alleles in this region are known to confer resistance to the fungus *M. oryzae* (Ma et al., 2022), whereas another encodes the MG1 immune receptor that recognizes RKN attack (Wang et al., 2023).

Long-term balancing selection provides an advantage for adaptation in plants by maintaining genetic diversity within a population or species. We hypothesize that long-term balancing selection is acting at the *MG1* and *KSL8* loci in *O. sativa* cultivars and contributes to maintaining such polymorphisms. Several studies in Brassica species have shown long-term balancing selection maintaining polymorphisms for immunity genes, such as NRLs, that span spatial and temporal scales (Bakker et al., 2006; Clark et al., 2007; Koenig et al., 2019; Kroymann et al., 2003; Wang et al., 2019; Wu et al., 2017). In *Boechera stricta*, an enrichment of NLRs was found within genomic regions under balancing selection (Wang et al., 2019). These regions are syntenic with *Arabidopsis thaliana* regions that also contain NLRs under balancing selection (Clark et al., 2007), suggesting maintenance of ancient NLR gene polymorphisms by balancing selection. Moreover, trans-species polymorphisms under balancing selection in *Arabidopsis* and *Capsella* are linked to genes related to biotic and abiotic stress responses (Wu et al., 2017).

The sessile nature of plants poses opportunities for dynamic and complex interactions with their environment, including synchronous and asynchronous temporal and spatial fluctuations in various biotic and abiotic factors. Such dynamic conditions and diffuse multi-species interactions can contribute to the maintenance of ancient polymorphisms in defense genes through balancing selection (Groen et al., 2016b; Karasov et al., 2014). It has been demonstrated in *B. stricta* that balancing selection is maintaining complex trait variation in the chemical profiles of leaves to withstand varying ecological factors, specifically herbivory and drought (Carley et al., 2021). It is well established that diterpenoids function to combat biotic and abiotic stresses (Schmelz et al., 2014; Zhao et al., 2018). As such, long-term balancing selection at the *KSL8* locus may contribute to maintaining polymorphisms and provide resilience against fluctuating combinations of stressors.

## Materials and Methods

### Plant material and growth conditions

Rice plants (*Oryza sativa* ssp. *japonica* cv. Kitaake, KitaakeX, Nipponbare, and knockout lines [Supplementary Table S12]) were grown under supplemental 16/8 hour light/dark conditions at 28°C in a greenhouse at the University of California Riverside’s Plant Research 1 facility. Rice seed preparation included a 3-day heat treatment at 50°C and sterilization (15% bleach for 5 minutes followed by sterile water washes). We germinated seeds on sterile wet paper for 6 days at 30°C in the dark. We transplanted seedlings into autoclaved soil composed of 70% sand and 30% organics. Seedlings were fertilized 4, 7, 10, and 14 days after transplanting with Peter’s solution (24-8-16) and additional Fe.

### Confirmation of rice knockout lines

To confirm the rice lines we are testing are true knockouts for their respective mutations we performed reverse transcription PCR (RT-PCR) (Supplementary Fig. S4,S5). For this analysis, root tissue was flash-frozen and ground to a fine powder. We isolated total RNA using the RNeasy Plant Mini Kit (Qiagen) following the manufacturer’s protocol. Residual DNA was removed using the TURBO DNA-*free* kit (Invitrogen) following the manufacturer’s protocol. From our RNA template, we produced cDNA using the High-Capacity cDNA Reverse Transcription Kit (Applied Biosystems) following the manufacturer’s protocol. We then performed RT-PCR to look for the presence of transcripts. *OsACTIN* was used as a control. Primers used in our analysis can be found in Supplementary Table S13.

### Root-knot nematode and fall armyworm husbandry

Root-knot nematodes of *Meloidogyne incognita* population “Project 77” (race 3) were cultured on tomatoes (*Solanum lycopersicum* cv. Moneymaker). To isolate RKN eggs, rice root systems were centrifuged for 3 minutes in a 10% bleach solution. From the suspension, eggs were collected from a 500-mesh sieve (Godinez-Vidal et al., 2024). RKN eggs were left for hatching on filter paper for 3 days in the dark at 24°C. We collected freshly hatched J2s for inoculum.

Second-instar fall armyworm (FAW) caterpillars (*Spodoptera frugiperda*) were purchased commercially (Benzon Research, Carlisle, PA).

### Co-expression enrichment analyses of herbivore-responsive genes

Herbivore-responsive genes were analyzed for enrichment in a rice co-expression network (Groen et al., 2022). We surveyed the differentially expressed genes (DEGs) identified from the following datasets: Kyndt et al. (2012), Zhou et al. (2020), Venu et al. (2010), Leclerc et al. (2024), Xue et al. (2024). RKN DEGs were assigned to root co-expression modules and FAW DEGs were assigned to shoot co-expression modules accordingly. We used a Fisher’s exact test designed for comparing count data to test for enrichment of DEGs within each module with the R package stats (R Core Team, 2022). We followed with a post-hoc test, the Benjamini-Hochberg test or false discovery rate (FDR) to adjust p-values for multiple comparison testing. Gene sets from significant modules underwent gene set enrichment analyses using ShinyGo v0.80 (Ge et al., 2020).

### Root herbivory assessment

For galling assays, individually potted 2-wk-old seedlings were inoculated with 350 J2s per seedling and maintained in the growth conditions described above. After 3 days and after 10 days, root growth of the rice knockout lines tested was similar to their respective wild-type and controls (Supplementary Fig. S6). At these time points, the rice plants were further evaluated for the number of RKNs present inside their roots, separating nematodes by life stage. For acid fuchsin staining, the roots were treated with 10% bleach for 5 min, washed thoroughly with water, and boiled for 10 s in acid fuchsin solution (3.5% acid fuchsin in 25% acetic acid). After letting the solution cool to room temperature, the roots were transferred to a de-staining solution (1 : 1 : 1, acetic acid : glycerol : H2O) before the number of nematodes that had entered the roots was evaluated using a dissecting microscope.

### Leaf herbivory assessment

Caterpillar weight gain assays were carried out as described previously (Groen et al., 2013). Individual second-instar caterpillars were transferred to leaves of each experimental plant using a fine paintbrush. The leaves with caterpillars were kept individually in plastic boxes with insect-proof lids and caterpillars were weighed to the nearest 0.1 mg using a micro-balance (Mettler-Toledo) before and after feeding for two continuous days.

## Data availability

Raw RNA sequence data from Groen et al. (2022) that some of our analyses rely on have been deposited as part of SRA BioProject PRJNA564338. Processed RNA expression counts, alongside a key to the RNA sequence data in SRA BioProject PRJNA564338 and the sample metadata were deposited in Zenodo under DOI 10.5281/zenodo.4779049. Normalized count data and further information on root- and shoot gene co-expression module composition and characteristics can be found in the supplemental material of Groen et al. (2022).

## Acknowledgments

This work was funded in part by the National Institute of General Medical Sciences of the National Institutes of Health (grant R35GM151194), the National Institute of Food and Agriculture (grant W5186), and University of California Riverside startup funds to S.C.G. We thank members of the Groen laboratory for helpful discussions.

## Disclosures

The authors have no conflicts of interest to declare.

## Notes

### Competing Interest Statement

The authors have declared no competing interest.

