## Supplementary Figures 1-6 for "Evolutionary Systems Biology Identifies Genetic Trade-offs In Rice Defense Against Above- and Belowground Attackers"

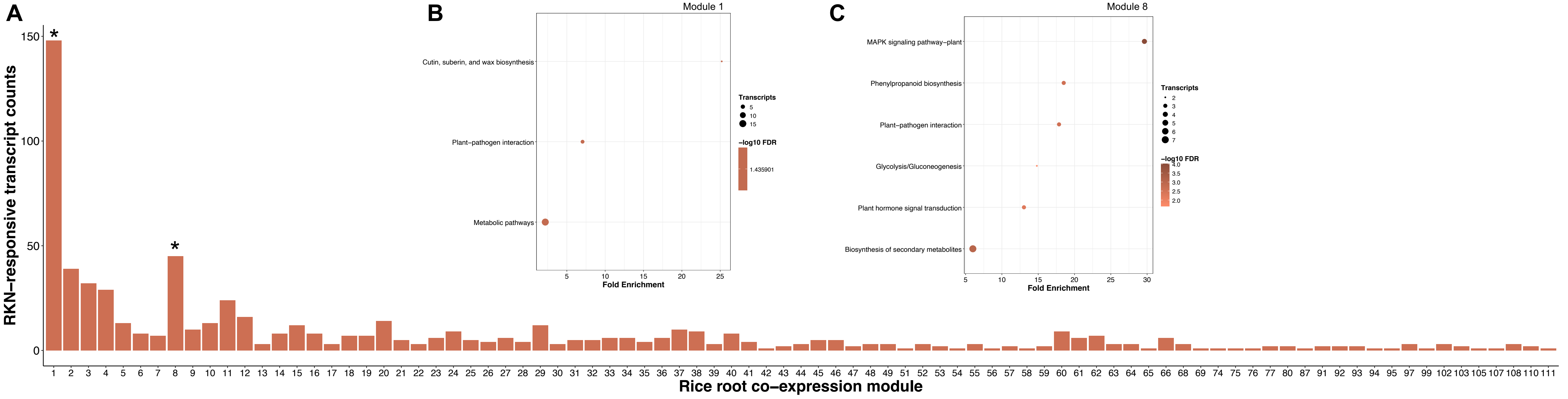


**Figure S1.** Distribution and gene set enrichment of RKN-responsive transcripts across the rice root co-expression network. A) Overlay of *Meloidogyne graminicola*-induced transcripts at 3- and 7-days post-infection combined on the rice root co-expression network. *, false discovery rate-adjusted p-value < 0.01 for a Fisher's exact test for count data. B,C) Enrichment of defense-related processes as shown by the KEGG enrichment analysis on root modules 1 and 8.

Dunivant et al., 2024

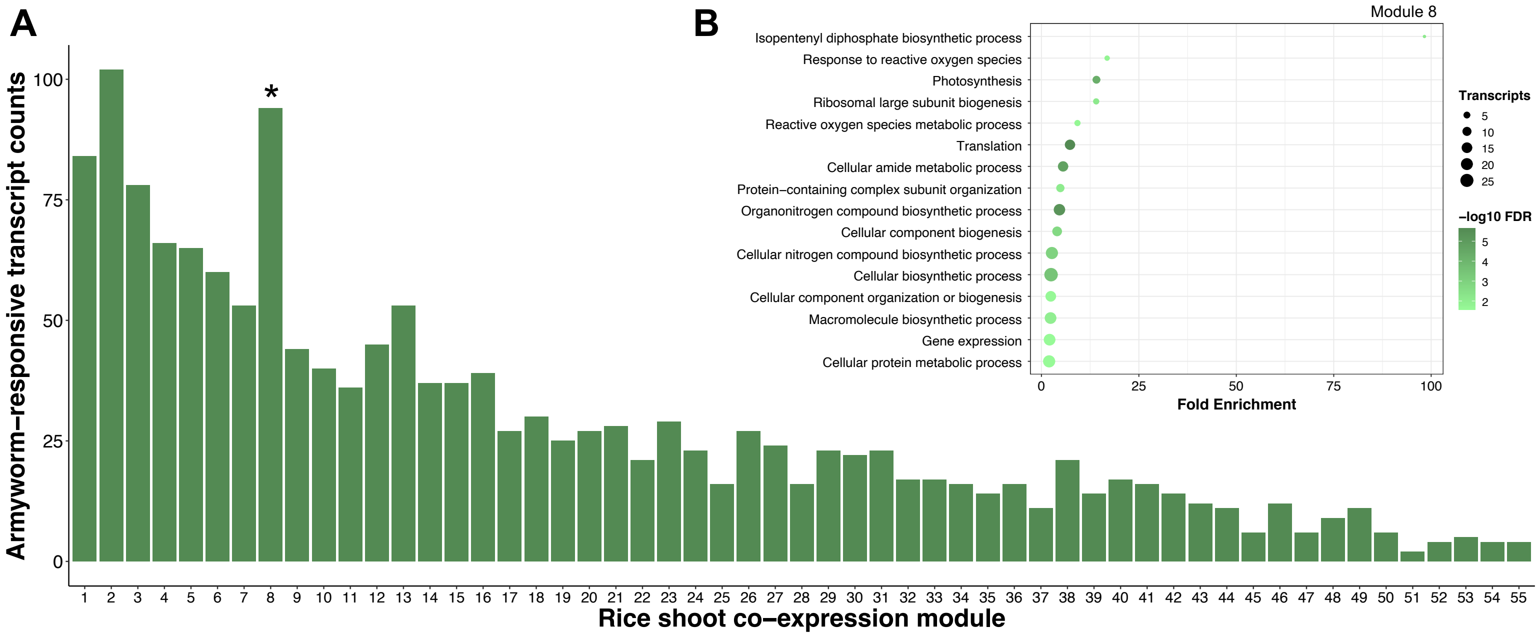


**Figure S2.** Distribution and gene set enrichment of armyworm-responsive transcripts across the rice shoot co-expression network. A) Overlay of *Spodoptera exigua*-induced transcripts at 1-day post-infection on the rice shoot co-expression network. *, false discovery rate-adjusted p-value < 0.01 for a Fisher's exact test for count data. B) Enrichment of defense-related processes as shown by the KEGG enrichment analysis of shoot module 8.

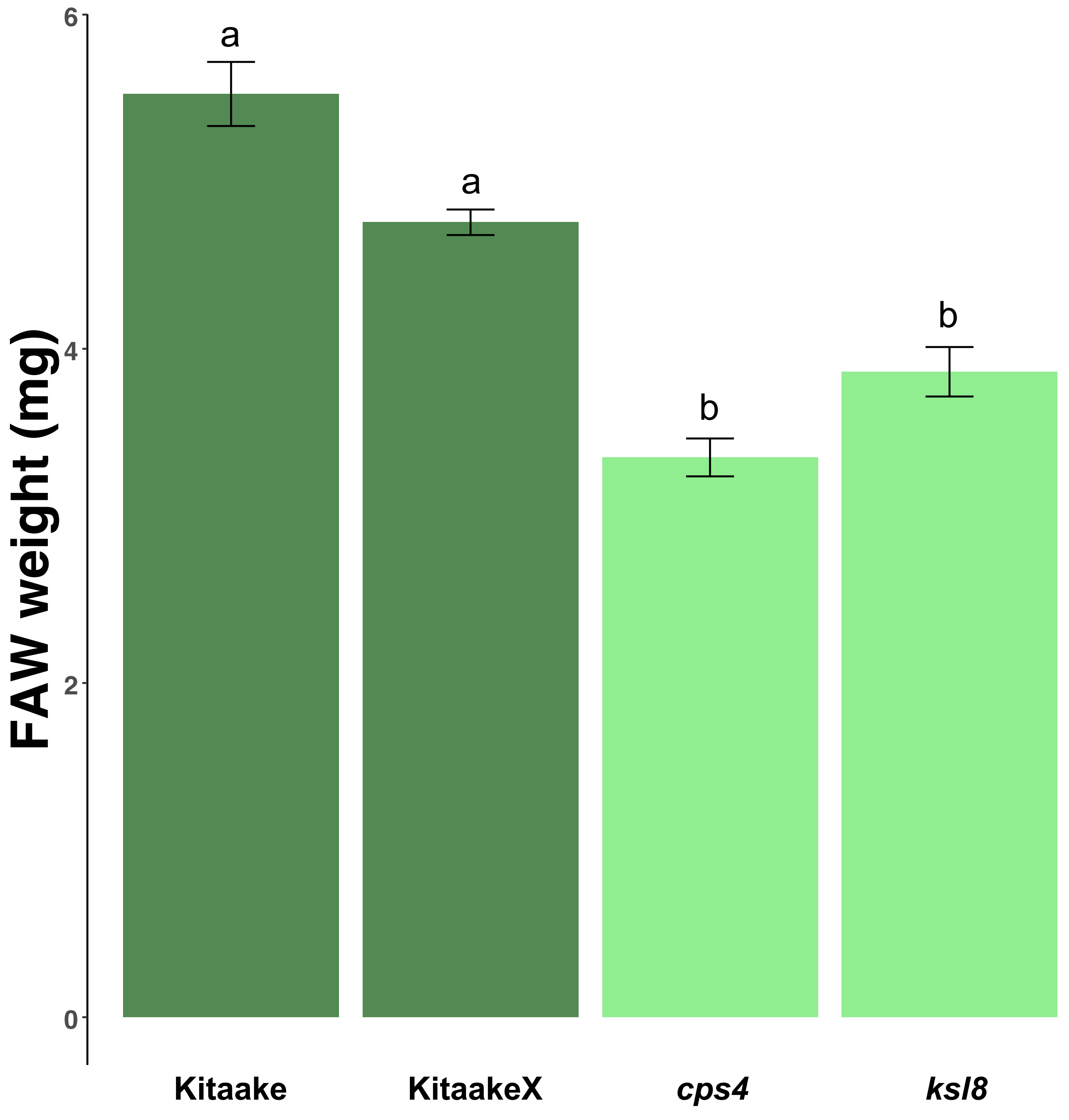


**Figure S3**. Changes in FAW weight after feeding for two continuous days on *cps4*, *ksl8*, and wild-type cv. Kitaake leaves.

**A** ***cps4*** **cv. Kitaake**

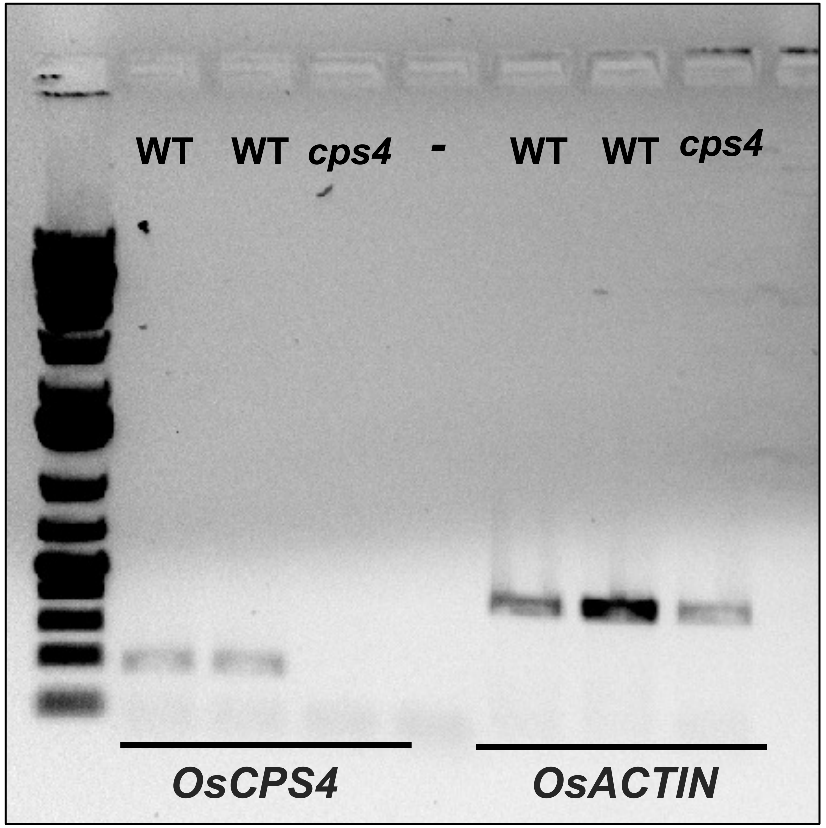


**B** ***ksl8*** **cv. Kitaake**

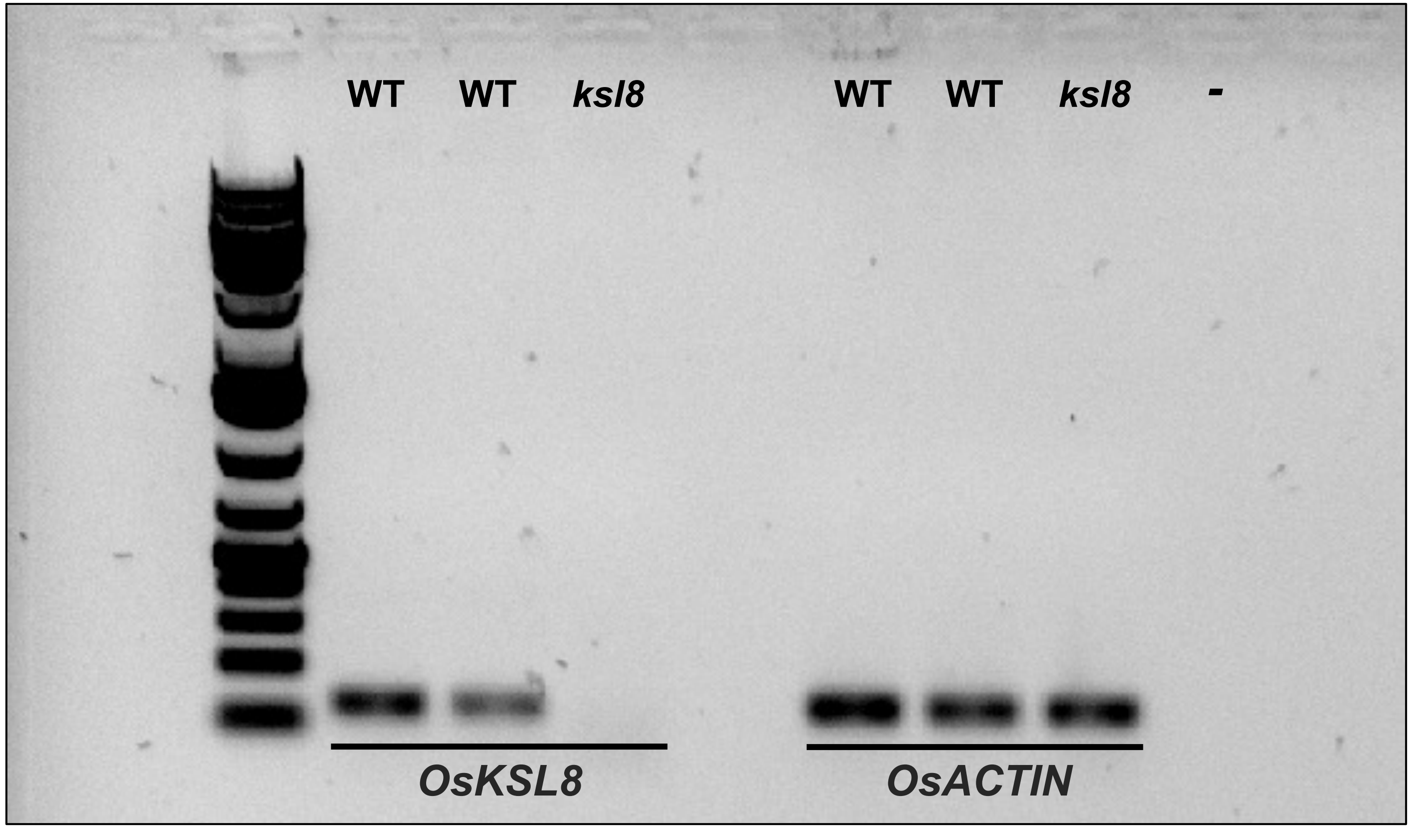


**Figure S4**. Gel image from RT-PCR to confirm *cps4* (A) and *ksl8* (B) cv. Kitaake knockout lines. *OsACTIN* was used as a control.

*cps4* cv. Nipponbare

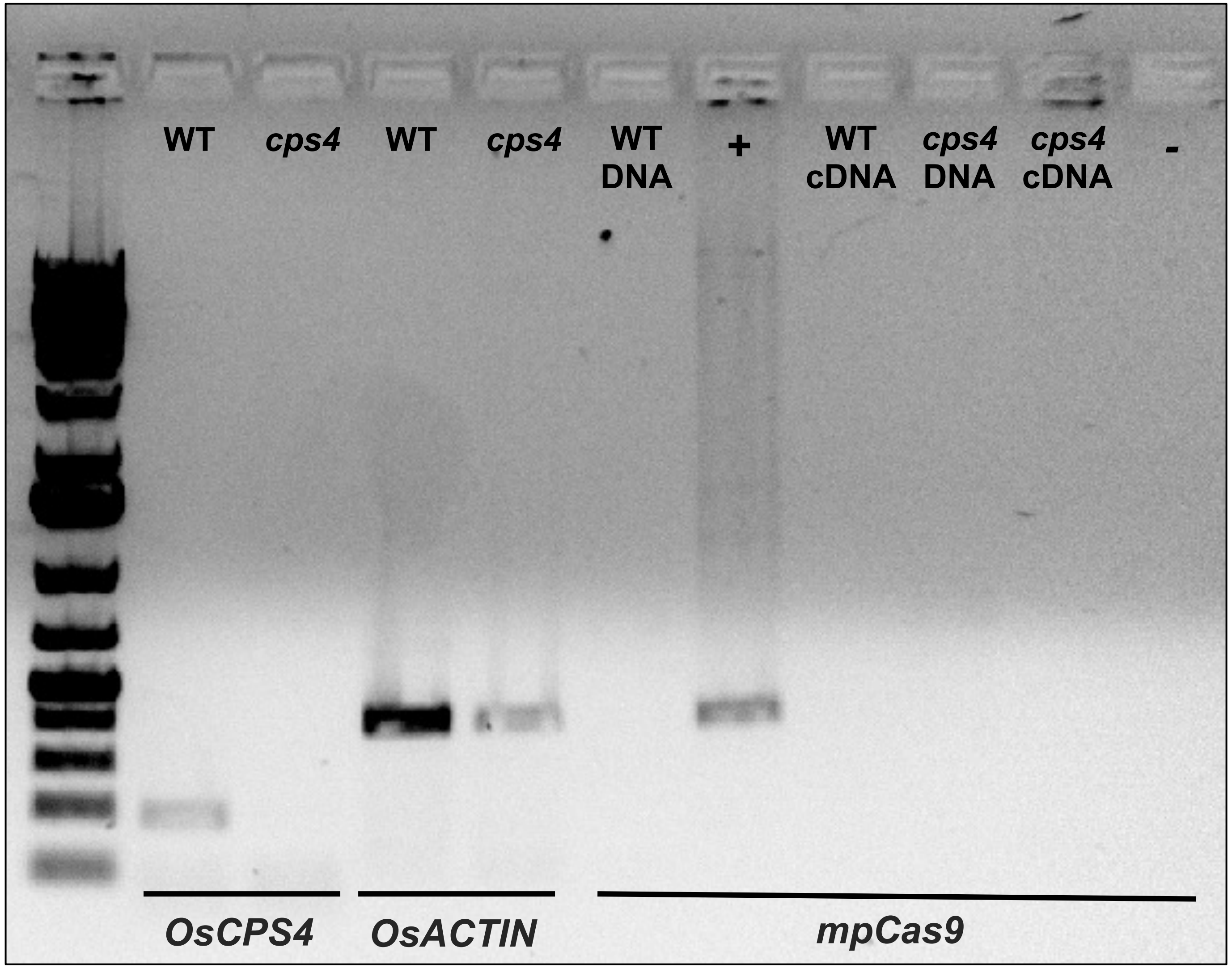


**Figure S5**. Gel image from RT-PCR to confirm *cps4* cv. Nipponbare knockout line. *OsACTIN* was used as a control. Additionally, *cps4* was confirmed to contain no residual Cas9 machinery.

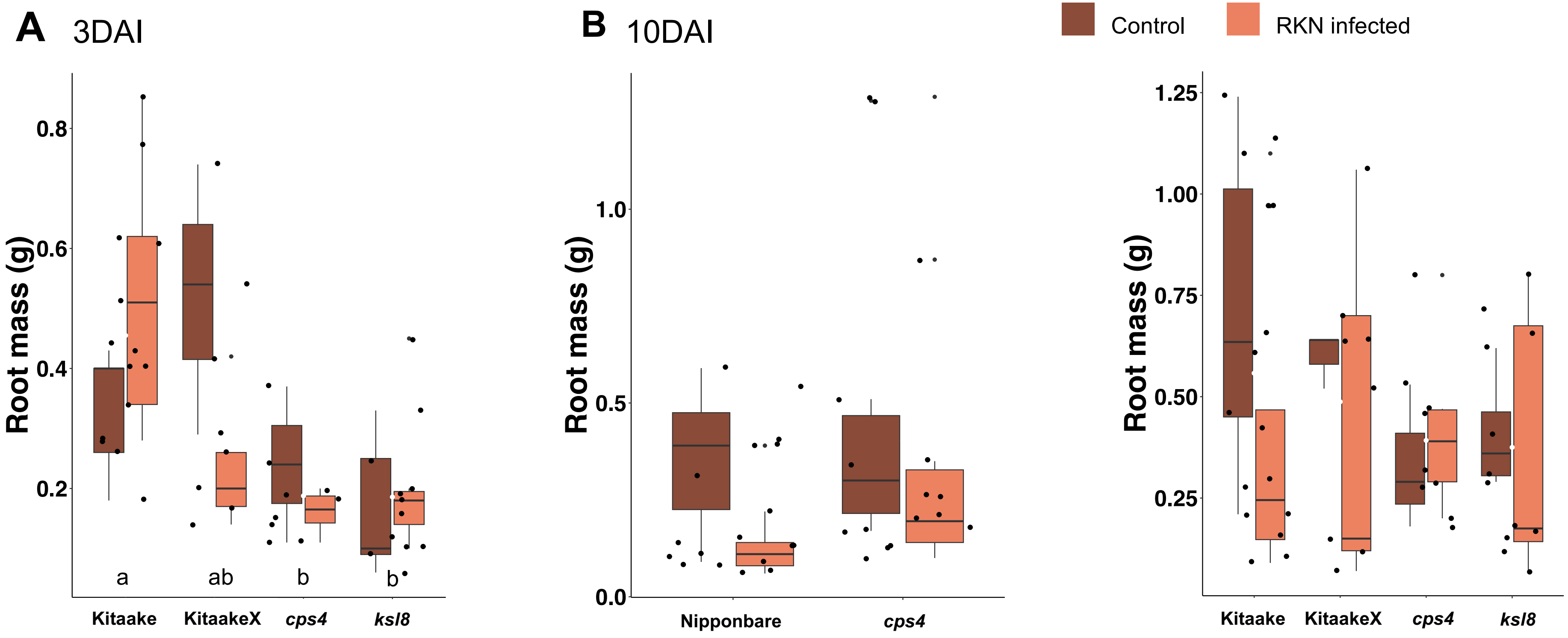


**Figure S6**. Final root mass of control and RKN-infected rice plants. A) *cps4*, *ksl8*, and wild-type cv. Kitaake plants for 3DAI. B) *cps4*, *ksl8*, and wild-type cv. Kitaake and Nipponbare plants 10DAI.

Dunivant et al., 2024
